# Virus-derived circular RNAs populate hepatitis C virus-infected cells

**DOI:** 10.1101/2023.07.28.551052

**Authors:** Qian Cao, Pakpoom Boonchuen, Tzu-Chun Chen, Shaohua Lei, Kunlaya Somboonwiwat, Peter Sarnow

**Affiliations:** Department of Microbiology & Immunology, Stanford University SOM, Stanford, CA 94305, USA; School of Biotechnology, Institute of Agricultural Technology, Suranaree University of Technology, Mueang, Nakhon Ratchasima, Thailand; Department of Biochemistry, Faculty of Science, Chulalongkorn University, Bangkok, Thailand; Natera, 201 Industrial Rd, San Carlos, CA 94070, USA; Center of Excellence for Leukemia Studies, St. Jude Children’s Research Hospital, Memphis, TN 38015, USA; Center for Applied Bioinformatics, St. Jude Children’s Research Hospital, Memphis, TN 38015, USA

**Keywords:** RNA viruses, Hepatitis C virus, circular RNA, viral internal ribosome entry site.

## Abstract

It is known that pre-mRNAs in eukaryotic cells can be processed to circular RNAs by a back-splicing mechanism. Circular RNAs have great stability and can sequester proteins or small RNAs to exert functions on cellular pathways. Because viruses often exploit host pathways, we explored whether the RNA genome of the cytoplasmic hepatitis C virus is processed to yield virus-derived circRNAs (vcircRNAs). Computational analyses of RNA-seq experiments predicted that the viral RNA genome is fragmented to generate hundreds of vcircRNAs. More than a dozen of them were experimentally verified by rolling-circle amplification. VcircRNAs that contained the viral internal ribosome entry site were found to be translated into novel proteins that displayed pro-viral functions. Furthermore, a highly abundant, non-translated vcircRNA was shown to enhance viral RNA abundance. These findings argue that novel vcircRNA molecules modulate viral amplification in cells infected by a cytoplasmic RNA virus.

**Significance Statement:** Processing of an RNA viral genome into hundreds of circular RNAs provides novel pro-viral functions and can promote translation of novel viral peptides.

## Introduction

It is well-established that pre-mRNAs in eukaryotic cells can be spliced into circular forms (circRNAs) by a nuclear back-splicing mechanism (1, 2). It has been estimated that approximately 25,000 circRNAs exist per cell at steady state (1, 3). These circRNAs are generated from an estimated 20% of transcribed genes (4), with hundreds of unique circRNAs being expressed in a tissue-specific manner. CircRNAs are more stable than their linearly spliced cousins due to their resistance to exoribonucleases (4). Very little is known about the biological functions for circRNAs. This is mainly because the abundance of most specific circRNA ranges only between 1 and 10 copies per cell (4). One of the most studied cytoplasmic circRNAs is CDR1as (or CirC7). CDR1as contains multiple binding sites for microRNA 7 and functions as a microRNA “sponge” to control midbrain development (5, 6). There is also accumulating evidence that circRNAs function in cell proliferation and immune regulation (7–9). Not surprisingly, certain DNA viruses, that use cellular pathways for transcription and RNA splicing, generate pre-mRNAs that are spliced into virus-derived circRNAs (vcircRNAs) (10). For example, Epstein-Barr virus encodes vcircRNAs that sequester microRNAs, resulting in the upregulating of genes that modulate viral malignancies (11, 12); Kaposi’s sarcoma virus generates vcircRNAs that modulate the lytic infectious cycle of the virus (13, 14); Human papillomavirus also generates a vcircRNA, E7, which is translated to encode the E7 oncoprotein (15). Here we demonstrate that the cytoplasmic hepatitis C virus (HCV) generates vcircRNAs which display pro-viral functions.

## Results

### Prediction of vcircRNAs generated from the HCV RNA genome

We conducted a study to identify any host circRNAs whose abundance was altered during HCV infection of hepatoma cells (16). Several of identified host circRNAs displayed pro-viral functions (16). Upon further interrogation of the RNase R-treated circRNA libraries used to identify these cellular circRNAs, we also identified, astonishingly, hundreds of putative vcircRNA species derived from the 10,000-nucleotide HCV genome (Table 1, Supplementary Table 1). A putative vcircRNA was defined by the presence of a discontinuous reading with two breakpoints in the junction region, and additional two nucleotides located at the start and end of junction-spanning reads (Figure 1A). Although most total viral reads were mapped to continuous regions on the viral genome, consistent with their templating from linear viral sequences, approximately 1% of the reads contained such novel junction sequences (Table 1). Most of these junction reads (97.7%) were derived from the positive strand of the RNA genome (Table 1). Although the 5’ and 3’ breaks points in each junction were distributed across the entire HCV genome, their locations were not random (Figure 1B). One cluster of reads (Cluster I) was observed at the 5’ end of HCV and contained both an intact viral internal ribosome entry site (IRES) and the sequences that encode the N-terminus of the viral core protein. The most abundant cluster (Cluster II) contained 5’ breakpoints between nucleotide positions 830 to 890 in the viral core coding sequence and 3’ breakpoints between 1150 to 1154 in the envelope E1 coding sequence (Figure 1B). In addition, another abundant cluster (Cluster III) was found within the NS5B region (Figure 1B). The most abundant vcircRNA junctions are shown in Figure 1C. The sizes of predicted vcircRNAs range from 130 nts to larger than 2000 nts, with an average size of 200-300 nts (Figure 1D). Analyses of the raw RNA-seq data using CIRI2 (https://sourceforge.net/projects/ciri/) also revealed predicted vcircRNAs with similar junctions, although the number of junction reads were much fewer (Supplementary Table 2).

**Figure 1.**
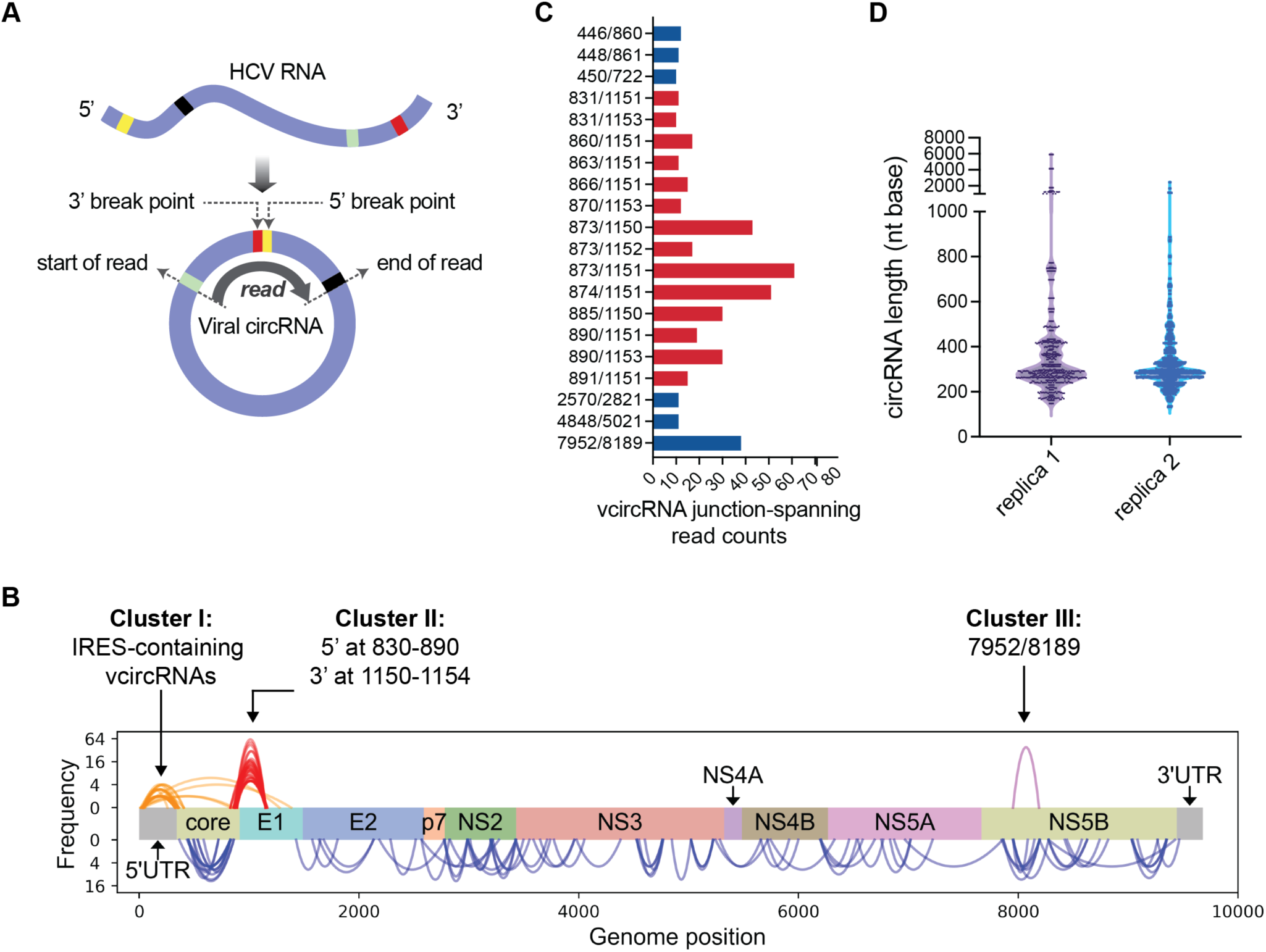
Landscape of HCV-derived circRNA (vcircRNA) junctions predicted by RNA-seq analysis. (A) Identification of putative vcircRNA junctions derived from the linear HCV RNA genome. The curved black arrow represents a junction-spanning read, which consists of two discontinuous sequences joined by their 5’ (yellow) and 3’ (red) break points. Additionally, the start position is depicted in mint green and the end position is shown in black when the read is aligned to the viral genome. To ensure the junction originates from vcircRNAs, it’s essential to validate the exact order of nucleotide positions, specifically the 5’ break point, end of read, start of read, and finally the 3’ breakpoint. (B) Distribution of predicted vcircRNA junctions on the entire HCV genome. Each junction read is indicated by an arch that connects its 5’ and 3’ break points. The three clusters examined in this study are indicated. (C) List of top 20 predicted vcircRNA junctions. (D) Length distribution of the vcircRNAs predicted from junction-spanning reads.

### Experimental verification of vcircRNAs

Although the presence of discontinuous junction sequences suggested their derivation from vcircRNAs, this needed to be experimentally verified. Reverse transcriptase-mediated rolling circle amplification was employed (17) to verify several members of the putative IRES-containing vcircRNA cluster I and the most abundant vcircRNA cluster II and III. Briefly, vcircRNAs were first reverse transcribed into cDNA products. These products, predicted to contain multiple iterated sequences of their template circular RNAs, were amplified by PCR using primers that flanked the novel junctions and, thus, diverged on linear viral counterpart sequences (Figure 2A). If template RNAs are truly circular, multiple rounds of transcription will occur, resulting in PCR products of various lengths that contain multiple junction sites, denoted as a, b, c, and d (Figure 2A). In contrast, linear RNAs should yield no products in the PCR reaction. Indeed, PCR fragments predictive of harboring multiple junction sequences were identified in infected, but not mock-infected cells (Figure 2B, Figure S1A). No specific PCR products containing junction sequences were amplified from full-length control viral RNAs that were *in vitro* transcribed by T7 RNA polymerase (Figure S1B), strengthening the argument that the vcircRNAs were generated in infected cells and did not represent PCR template switching or other artifacts. Sanger sequencing of the cloned PCR products revealed complete, iterated sequences of vcircRNAs in all the three clusters of vcircRNAs (Figure 2C). Each iterative sequence was flanked by the predictive identical junction sites. So far, 14 out of 14 predicted vcircRNAs have been verified by rolling circle RT-PCR and Sanger sequencing (Supplementary Table 3).

**Figure 2.**
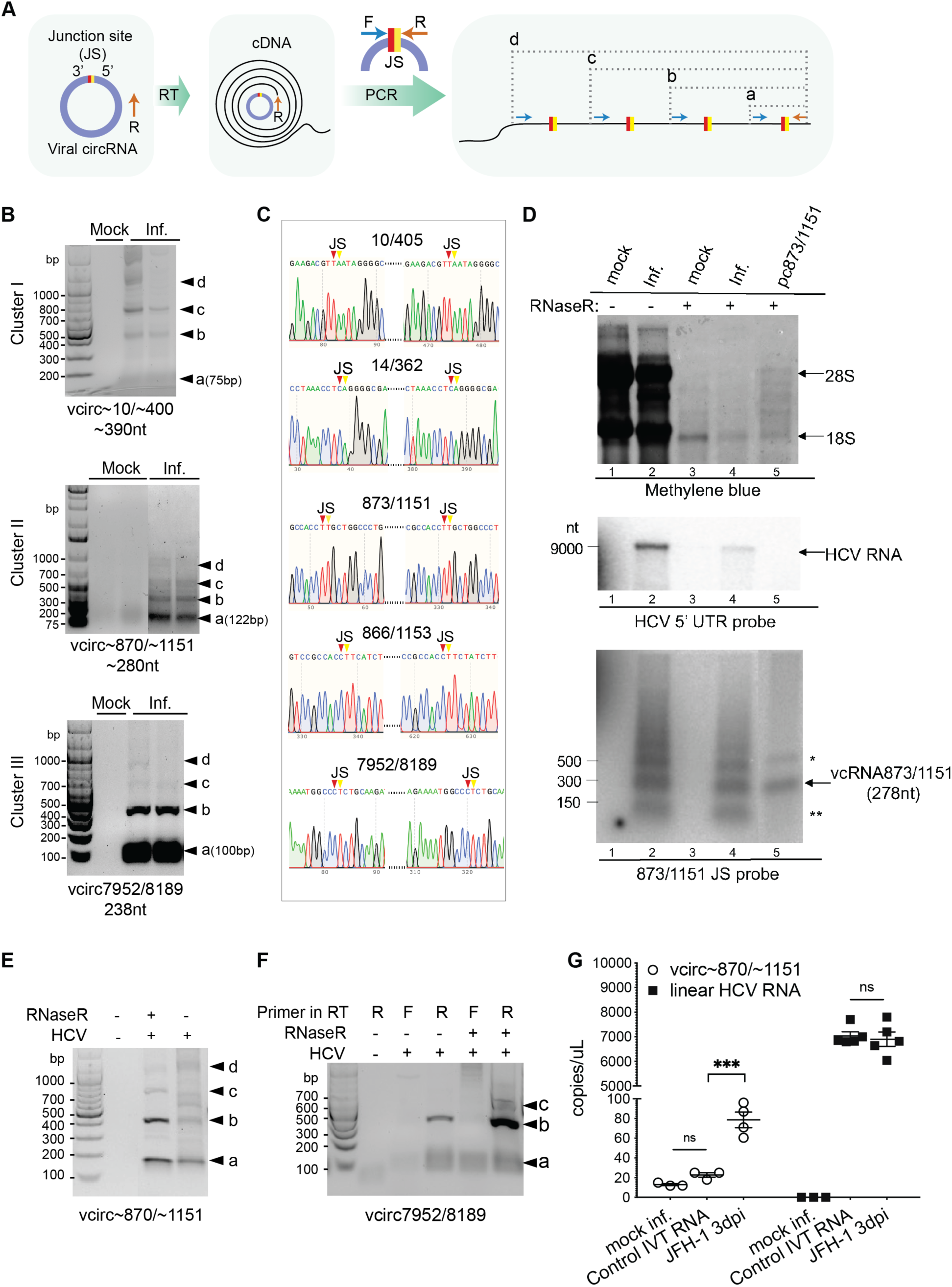
Identification of HCV vcircRNAs in HCV infected cultured hepatoma cells. (A) Schematic views of vcircRNAs subjected to reverse transcription and subsequent PCR using primers that flank the junction sites of predicted vcircRNAs but diverge on linear RNAs. RT-PCR products of single and multiple repeats of the vcircRNA sequences are denoted as b, c, and d while the smallest amplicon “a” only contains a junction sequence (right panel). (B) Agarose gel electrophoresis of RT-PCR products. PCR products representing multiple copies of vcircRNAs from cluster I (IRES-containing vcircRNA∼10/∼400), cluster II (the most abundant vcircRNA∼870/∼1151) and cluster III (vcircRNA 7952/8189) are shown. Primers flanking the variable junction sites of cluster I or cluster II vcircRNAs were used. For cluster III 7952/8189, junction-spanning primers were used (listed in Supplementary Table 4). Nested PCR was performed to amplify cluster I vcircRNAs due to their low abundance in the infected cells. Shown here (upper panel in B) is a representative image of the 2^nd^ round PCR for cluster I vcircRNAs. (C) Representative Sanger sequencing results of RT-PCR products inserted into a TOPO vector. Each sequence trace displays two junction sites denoted as triangles with one full copy of the indicated vcircRNA sequence. (D) Northern blot detection of cluster II vcircRNA 873/1151 from HCV JFH-1-infected cells. Total RNA was extracted from uninfected or infected Huh7 cells at 3 d.p.i. Lane “pc873/1151” denotes RNA isolated from plasmid-based expressed circRNA873/1151 (see also Figures S2A-S2C). Ribosomal RNA 28S and 18S are indicated by methylene blue staining (top panel) of the membrane. A 5’ UTR probe (84-374nt) was hybridized to detect HCV genomic RNA (middle panel). An oligonucleotide probe specific for the junction site was hybridized to detect vcircRNA 873/1151 (bottom panel). * and ** suggests the presence of RNAs with different topologies or of broken RNAs. (E) RT-PCR amplification of cluster II vcircRNAs with or without RNase R treatment. (F) Strand-specificity and RNase R resistance of cluster III vcircRNAs. After RNase R treatment (30 min at 37 °C), RNA was reverse transcribed with a specific reverse primer or forward primer spanning the junction (listed in Supplemental Table 4) to differentiate between plus-stranded or negative-stranded vcirc7952/8189 followed by subsequent PCR. (G) Copy numbers of cluster II vcircRNAs quantified by droplet digital PCR. Divergent primers that flank the junctions of ∼870/∼1151 were used to amplify the vcircRNA (open circles), while convergent primers targeting the HCV Core region (squares) were used to amplify linear viral RNA. “Control IVT RNA” is *in-vitro* transcribed full-length HCV RNA. Error bars indicate means ± SEM. Statistical significance was determined by ordinary one-way ANOVA. ns, not significant, ***, p<0.001.

To further validate the presence of viral circRNAs, Northern blot analysis using an oligonucleotide probe with sequence complementarity to the junction site of vcircRNA 873/1151 was performed. Extensive degradation of host ribosomal RNA (Figure 2D, upper panel) and HCV genomic RNA (Figure 2D, middle panel) was observed in RNA samples treated with RNase R, which selectively digests linear RNA. In contrast, vcircRNA ∼873/∼1151 was detected in nuclease-treated samples obtained from vcircRNA-overexpressing cells (pc873/1151; Figures S2A-2C) and infected cells (Figure 2D, bottom panel). Finally, rolling circle RT-PCR amplification also revealed the presence of cluster II (Figure 2E) and mostly positive-stranded cluster III (Figure 2F) vcircRNAs in the RNase R-treated samples. Thus, the junction-specific RT-PCR, Northern blot analysis and RNase R results argue that *bona fide* vcircRNAs populate HCV-infected cells. Droplet digital PCR (ddPCR) showed that the abundance of vcircRNA∼870/∼1151 is approximately 1% of HCV genomic RNA in infected cells (Figure 2G). In control experiments using *in vitro* transcribed full-length viral RNA, linear HCV RNAs can be easily detected by ddPCR, while only background levels of vcircRNAs could be observed (Figure 2G). These findings strongly argue that this vcircRNA is a genuine product of viral infection.

### Translation of cluster I vcircRNAs

In 1995, we showed that synthetic circRNAs that contained the IRES of encephalomyocarditis virus (EMCV) could be translated (18). Thus, we hypothesized that the HCV IRES-containing vcircRNAs represent a class of naturally derived vcircRNAs that can be translated into novel proteins. We determined the abundance of cluster I vcircRNAs to be approximately 0.25% of the abundance of linear viral RNA (Figure S2D) and verified the circularity of several members of the cluster I vcircRNAs (vcircRNA10/405, vcircRNA 26/369, vcircRNA30/372 and vcircRNA14/362) by rolling circle RT-PCR and Sanger sequencing. To determine whether these HCV IRES-containing circRNAs could mediate translation, they were expressed from a split-GFP plasmid, in which the C-terminal coding sequence of GFP is placed before its N-terminal sequence, separated by the individual vcircRNA sequence that contain the HCV IRES (Figure S2E). This cassette was flanked by two inverted *Drosophila lacasse2* intronic repeats, which mediate back-splicing events and facilitate the circularization of the linear transcript, thus allowing IRES-mediated production of full-length circular GFP in transfected cells (19). Control EMCV IRES-containing circRNAs produced GFP efficiently in transfected cells (Figure 3A). In contrast, the 873/1151 HCV sequence, which does not contain an IRES, did not promote the synthesis of GFP (Figure 3A). Gratifyingly, all IRES-containing HCV sequences expressed GFP (Figure 3A), demonstrating that the HCV IRES is active when placed into a circular RNA.

**Figure 3.**
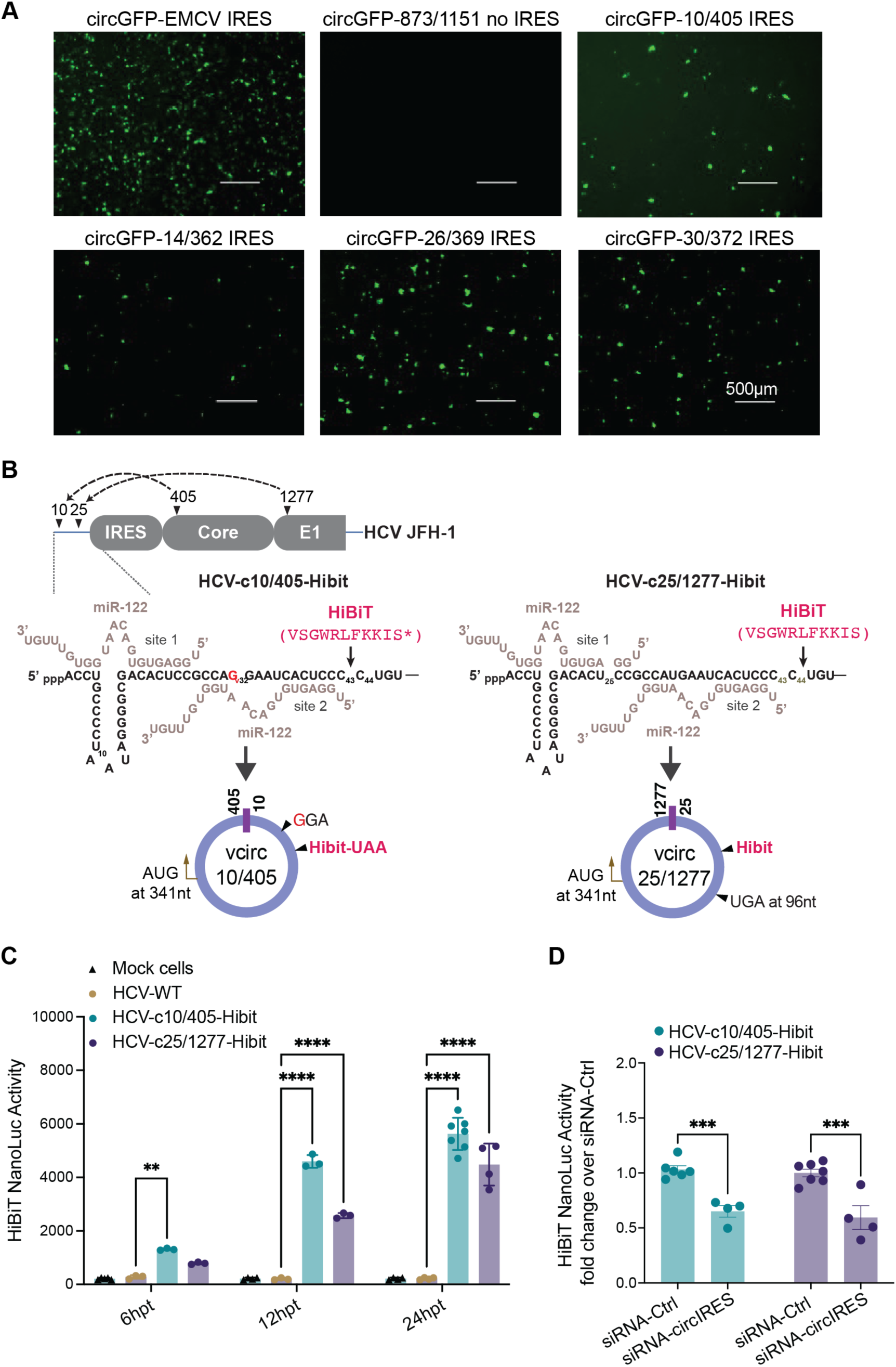
Translation of cluster I IRES-containing vcircRNAs. (A) Translation of the HCV IRES within a circular GFP RNA expressed from a polymerase II promoter-containing split-GFP plasmid that promotes back-splicing to create circular RNAs (see also Figure S2E). Plasmids contain the IRES from encephalomyocarditis virus (EMCV), or individual vcircRNA sequence were transfected into Huh7 cells for 48 hours, and GFP fluorescence was visualized. (B) Illustrations of two infectious clones of HCV that contain a HiBiT tag of 11 amino acids(20) inserted into the 5’ noncoding region of HCV after the microRNA 122 binding sites, so as not to interfere with microRNA 122 binding (21). Synthesis of the HiBiT tag from the natural AUG codon at HCV 341nt will demonstrate that HiBiT-containing cluster I circular RNAs are generated and translated. The HCV-c10/405-Hibit mutant has an additional U to G mutation (the G is highlighted in red) to eliminate the upstream UGA codon and allow the expression of the HiBiT tag at the C-terminus of vcirc10/405 ORF. In the HCV-c25/1277-Hibit mutant, HiBiT is inserted in-frame with the vcirc25/1277 ORF. (C) HiBiT expression from infectious viral genomes described in Figure 3B. HiBiT expression from wildtype HCV or mutants were measured as HiBiT NanoLuc activities at indicated time points. (D) Effects of junction-specific siRNAs on HiBiT expression from IRES-containing vcircRNAs in virus-infected cells. Huh7 cells were transfected with control siRNA (siRNA-Ctrl), or a pool of siRNAs targeting the junction sites of circ10/405, circ26/369, circ30/372 and circ25/1277 at a final concentration of 100 nM. At one day post transfection, cells were transfected with full-length viral RNA of HCV-c10/405-Hibit or HCV-c25/1277-Hibit. HiBiT expression was analyzed at 24 hours after viral genome transfection. At least three replicas were used for each group. Error bars indicate means ± SEM. Statistical significance was determined by ordinary two-way ANOVA. *p<0.05, **p<0.01, ***p<0.001, ****p<0.0001.

Although the IRES-containing vcircRNAs are of various lengths, all of them contain nucleotides 42-356 and are predicted to use the AUG at nucleotide 341 that is normally used for translation initiation of the HCV core protein. The proteins encoded by these vcircRNAs, depending on the location of their junction sequences, are predicted to contain a partial sequence of HCV core protein, followed by a novel sequence encoded from their 5’ break points in the junction site until a stop codon is met. To determine whether the IRES-containing vcircRNAs were translated when produced naturally from the HCV genome in infected cells was challenging, as appropriate antibodies to the novel products are not yet available. We chose two different cluster I vcircRNAs, 10/405 and 25/1277, to monitor the production of 29 and 336 amino acid proteins respectively (Figure S3). To enable their detection, a sequence encoding a small peptide tag of 11 amino acids, called HiBiT (20), was inserted downstream of the 5’ junction site of vcircRNA 10/405 and 25/1277 in the non-coding region of HCV genome (Figure 3B). Care was taken not to disrupt the two microRNA122 binding sites which are essential for viral gene amplification (21). The HiBiT sequence has been engineered to be a required component of the luminescent NanoBiT enzyme, allowing HiBiT-containing proteins to be quantified as bio-luminescence (20, 22). Since it is placed in the 5’ UTR of HCV genome, HiBiT is exclusively expressed from IRES-containing vcircRNAs. Two variant genomes were constructed to test the translatability of the IRES-containing vcircRNAs (Figure 3B): HCV- c10/405-Hibit, which is predicted to generate a novel core-HiBiT fusion peptide, and HCV- c25/1277-Hibit, which is predicted to generate a HiBiT fusion protein from a different reading frame (Figure S3). Both wildtype and variant RNA genomes were expressed in transfected cells (Figure S4). Next, translation of the IRES-containing vcircRNAs was monitored. Unlike the HiBiT-lacking wild-type HCV, transfection of either the HCV-c10/405-Hibit or HCV-c25/1277- Hibit RNA genomes gave rise to significant luminescence (Figure 3C). This result indicates that both vcirc10/405 and vcirc25/1277 can be translated when expressed from full-length viral RNA during viral infection. In further support that the observed luminescence resulted from translation of vcirc10/405-Hibit and vcirc25/1277-Hibit, siRNAs directed to the junction sequences of IRES- vcircRNAs were tested and shown to diminish HiBiT accumulation (Figure 3D). To exclude the possibility that HiBiT was translated from linear RNA in an IRES-independent manner, linear control RNA fragments from the 5’ end of HCV-c10/405-Hibit were generated, and their translation were examined after transfection. Compared to the full-length HCV-c10/405-Hibit RNA, all HiBiT-containing linear control RNAs yielded only background luminescence (Figures 4A-4B), strongly arguing that HiBiT expression originated exclusively from vcircRNAs. These experiments support the conclusion that IRES-containing vcircRNAs can be translated into novel products in HCV-infected cells.

**Figure 4.**
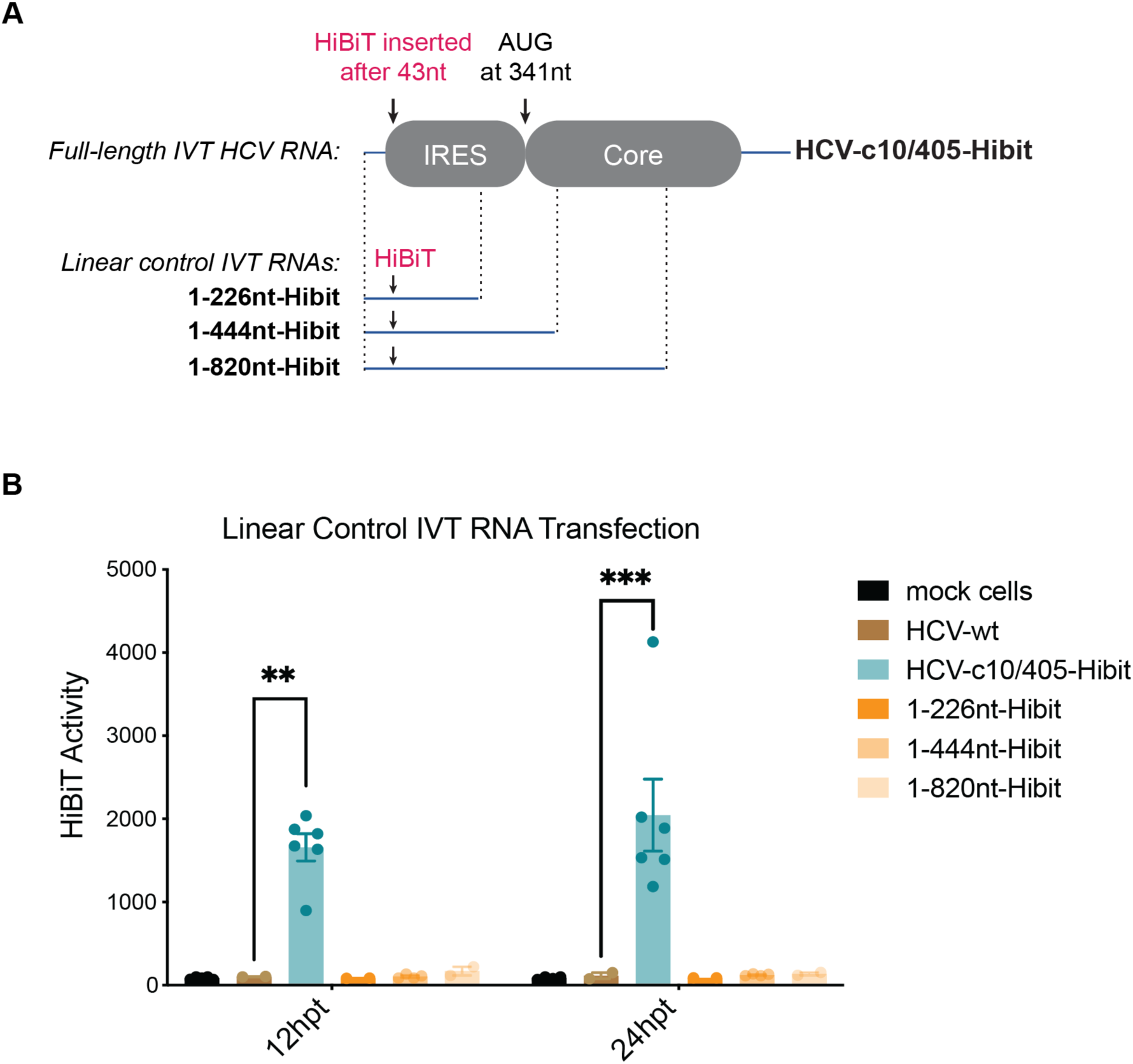
Translation of the HiBiT-containing linear control IVT RNAs. (A) Schematic view of designing short linear control RNAs of various lengths derived from HCV-c10/405-Hibit. Linear RNA fragments containing HiBiT were generated by *in vitro* transcription of indicated PCR products amplified from 5’ end of the mutant plasmid of HCV-c10/405-Hibit. (B) Huh7 cells in 12-well plates were transfected with 1 μg of each linear control RNA or *in vitro* transcribed full-length HCV-WT (lacking HiBiT) or HCV-c10/405-Hibit RNAs. At 12 or 24 after transfection, HiBiT expression was quantified in the HiBiT Lytic Assay (Promega). At least four replicas were used for each group. Error bars indicate mean ± SEM. Statistical significance was determined by ordinary two-way ANOVA. **p<0.01, ***p<0.001.

### Functions of vcircRNAs during viral infection

Do the identified vcircRNAs have pro- or anti-viral functions in HCV-infected cells? To test this, vcircRNAs were depleted by specific siRNAs in cells either infected with HCV or transfected with full-length wild-type HCV RNA. Figure 5A shows that depletion of either IRES-containing cluster I vcircRNAs and cluster II vcircRNA 873/1151 resulted in a significant decrease in viral RNA abundance in both virus-infected and viral RNA-transfected cells. To test for potential off-target effects of siRNAs on linear HCV RNAs, which might reduce their abundance, a replication-deficient HCV JFH-1 genome carrying a Nanoluciferase gene (HCV-Nluc-GND) was generated (Figures S5A-5C). Transfection of vcircRNA-targeting siRNAs did not alter the Nano- luciferase expression produced from the replication-deficient HCV (Figure 5B), arguing that the siRNAs do not display off-target effects on linear viral RNA sequences, and that both IRES-containing vcircRNAs and vcircRNA 873/1151 have pro-viral functions.

**Figure 5.**
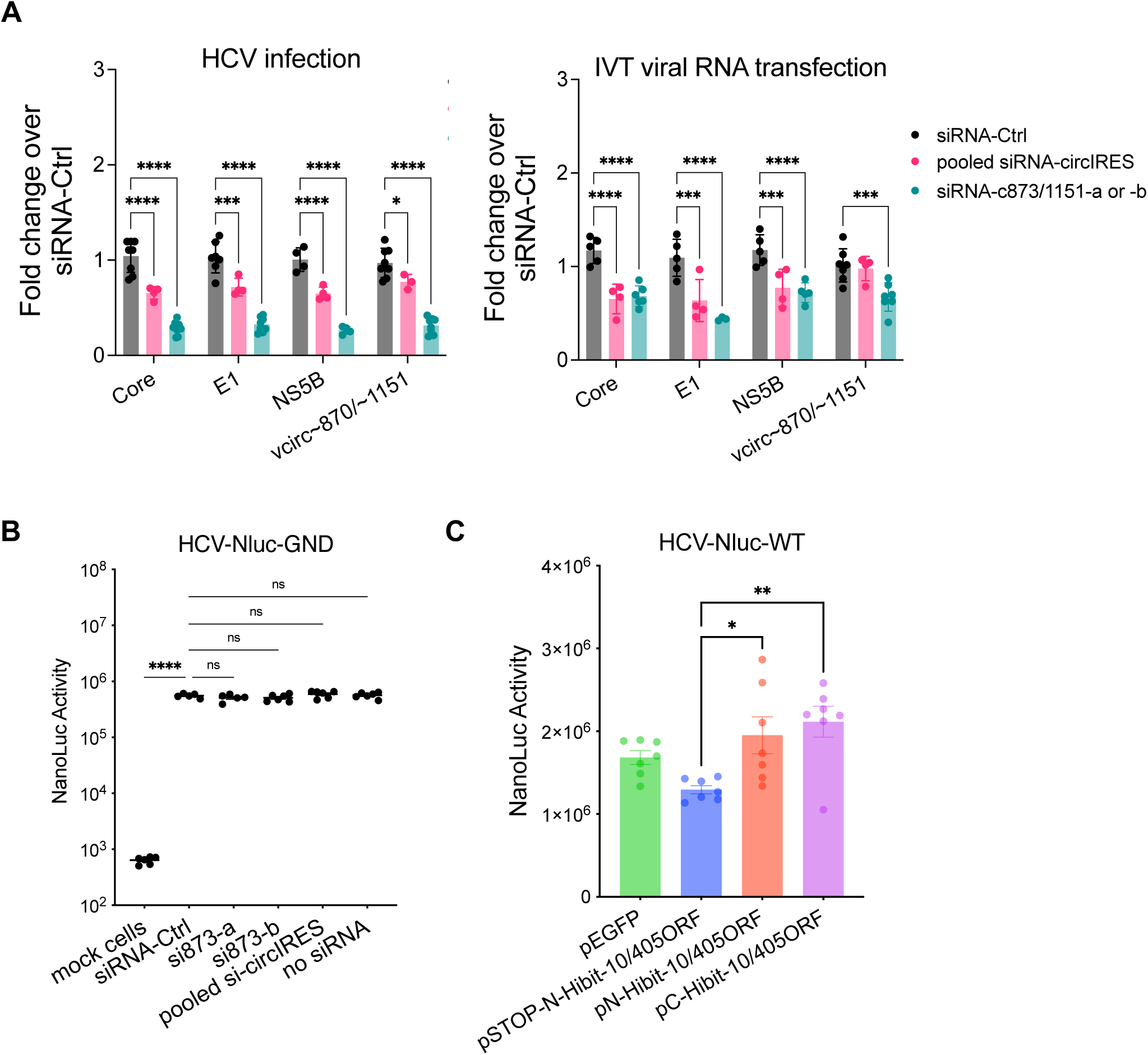
Effects of vcircRNAs and the translational product of cluster I vcircRNA on HCV infection. (A) Effects of cluster I and II vcircRNA depletion on viral RNA abundances in virus- infected or RNA-transfected cells. Huh7 cells were transfected with control siRNA, pooled siRNAs against IRES-containing circRNAs, or siRNA-a or -b targeting the junction site of non- IRES vcirc873/1151 at a final concentration of 100 nM. At one day-post-transfection, cells were further infected with HCV JFH-1 strain at a multiplicity of infection of 0.5 (left panel) or transfected with full-length IVT wild-type viral RNA (right panel). Two days later, HCV RNA abundance at Core, Envelope 1, NS5B regions of the linear RNA genome, as well as the junction sequences in vcirc∼870/∼1151 were quantified by qRT-PCR. At least four biological replicates were used for each group. (B) Effects of siRNAs on translation of a replication- defective HCV RNA containing a luciferase reporter (HCV-Nluc-GND). Huh7 cells were transfected with indicated siRNAs for 24 hours, and then transfected with *in vitro* transcribed HCV-Nluc-GND RNA. HCV Nluc activity was further measured by NanoGlo assay 20 hours later. (C) Effects of vcircRNA10/405 ORF overexpression on HCV reporter virus (HCV-Nluc- WT). Huh7.5.1 cells were transfected *in vitro* transcribed HCV-Nluc-WT RNA for 24 hours, and then transfected with each of control EGFP, control Stop codon-ORF, N-terminal or C-terminal tagged ORF plasmids. After 24 hours HCV Nluc activities were measured in NanoGlo assays. At least four replicas were used for each group. Error bars indicate means ± SEM. Statistical significance was determined by ordinary two-way ANOVA. ns, not significant, *p<0.05, **p<0.01, ***p<0.001, ****p<0.0001.

Finally, it was examined whether the protein product encoded by the IRES-containing vcircRNA10/405 enhanced viral gene expression. Overexpression of N- or C-terminal HiBiT-tagged vcircRNA10/405 open reading frames, but not a control reading frame interrupted by two stop codons (pSTOP-N-Hibit-10/405ORF), enhanced viral gene expression (Figure 5C, Figures S6A-S6B), arguing that the 29-amino acid peptide itself has a pro-viral function. These results indicate that novel peptides generated by IRES-containing vcircRNAs, and vcircRNAs themselves can enhance viral RNA abundances, likely by distinct mechanisms.

## Discussion

This study shows that vcircRNAs with pro-viral functions are generated from distinct parts of the HCV RNA genome, thus presenting a novel class of viral RNA species that populate infected cells. Some of HCV-derived vcircRNAs contain the HCV IRES, and their unique open reading frames are translated. It is unlikely that the non-IRES-containing vcircRNAs all have distinct functions, considering that hundreds of them were predicted in our RNA-Seq study. Abundant vcircRNAs could sequester proteins or small RNAs that modulate viral infection, or they could function in bulk after translocating to uninfected bystander cells (23). Perhaps vcircRNAs inhibit initial innate immune responses, for example by inhibiting the activation of protein kinase R in uninfected bystander cells (24). A limitation of this study is that it was performed in cultured liver cells that were infected with the HCV JFH1 genotype 2 which replicates efficiently in Huh7 cells. While animal (25) and organoid models (doi.org/10.1101/2021.10.26.465357) for HCV are available, they are not very robust to study viral gene expression. However, putative vcircRNAs have been detected in the silkworm midguts infected with *Bombyx mori* cypovirus, a member of the reoviridae (26). Thus, it likely that HCV vcircRNAs are produced in animal tissues as well.

At present, it is not clear by what mechanism these vcircRNAs are being generated from HCV whose infectious cycle resides exclusively in the cytoplasm. There is no evidence that the cellular splicing machinery is re-located to the cytoplasm in infected cells, and HCV viral RNA cannot be detected in the nucleus (27). Furthermore, sequence alignment in the proximity of the junction sites did not yield any conserved motifs commonly seen in host pre-mRNA splicing, suggesting that these RNA virus-derived circular RNAs are probably generated by pathway(s) that are distinct from those utilized by cellular circRNAs involving splicesomes (28). We speculate that a genome-wide knockout screen study could provide valuable insights into host factors responsible for the biogenesis of HCV circRNAs.

There is evidence that other RNA viruses, such as SARS coronaviruses (29–31) and the *Bombyx mori* reovirus (26, 32) accumulate RNA species that contain non-contiguous viral sequences. In those studies, selected junction sequences were demonstrated by RT-PCR. However, it cannot be ruled out that some of these RNAs represent linear recombinant RNAs. Here we validated complete sequences of vcircRNAs by rolling circle amplification and demonstrated their presence during infection by Northern blot analyses with resistance to RNAseR. Future investigations of novel vcircRNAs across a diverse range of RNA viruses will reveal the prevalence of virus-derived circRNAs and their potential significance in viral growth and pathogenesis. Such studies will ultimately contribute to the development of effective therapeutic strategies with which these novel virus-derived RNAs can be targeted.

## Methods

### Cell Culture, HCV infection and RNA extraction

The human hepatocarcinoma cell lines Huh7 and Huh7.5.1 were maintained in Dulbecco’s modified Eagle’s medium (DMEM, Invitrogen) supplemented with 10% heat inactivated fetal bovine serum (FBS), pen/strep and non-essential amino acids. Cells were grown in an incubator with 5% CO2 at 37 °C. For infection, six well-plates were seeded with 2.5×10^5^ Huh7 cells per well. On the next day, supernatants were removed, and cells were infected with HCV JFH-1 viral stock at MOI of 0.1 for 3 hours. The inoculum was removed and replaced with 2 ml of DMEM. At 48 or 72 hours-post infection, total RNA from infected and non-infected cells was extracted using TRIzol reagent (Invitrogen), following the manufacturer’s instructions. Genomic DNA was removed in-column by Turbo DNase I treatment (ThermoFisher) using RNA concentrate and clean kit (ZYMO lnc.).

### Sample preparation for RNA-Seq

RNA samples with an RNA integrity ≥ 9 were first rRNA-depleted, and then treated with RNase R (Lucigen). After phenol/chloroform/isopropanol extraction, samples were used to construct RNA-Seq libraries, using the True-Seq Stranded Total RNA Library Kit (Illumina Inc., USA).

These libraries were then sequenced on the Illumina sequencing platform (NovaSeq 6000 platform) and 150 bp/125 bp paired-end reads were generated. Illumina sequencing and QC of the raw data were performed at Stanford Genomics (Stanford, CA, USA).

### *In silico* identification of viral circular RNAs (vcircRNA)

The Galaxy tool (https://usegalaxy.org/) was used to perform Q30 and adaptor trimming on the raw data, and the reads were aligned with the JFH-1 reference genome (GenBank No. AB047639). An expectation value cutoff of 10^-5^ was used in the blastn. On the blast-hit table, those reads that mapped to two discontinuous regions on the viral genome were further subjected to an analysis pipeline searching for the vcircRNA junction-specific reads, according to the correct order of four nucleotide positions in the discontinuous mappings when reads are aligned to the viral genome.

### Validation of HCV vcircRNAs using rolling circle amplification and PCR

1μg RNA was reverse transcribed using the Superscript III kit (Invitrogen) as described in the manufacturer’s instruction. A reverse primer specific to the predicted circRNA was added to the reaction to increase the yield of cDNA from circRNAs (Supplementary Table 4). Subsequently, PCR (PrimeSTAR MAX DNA polymerase, Takara Bio USA, Inc.) was performed using divergent primers flanking or spanning the junction site of the specific circRNA. The oligonucleotides used in this study are listed in Supplementary Table 4. The cycling condition was as follows: 98°C 30s, 34 cycles of 98°C 10sec, 59°C 5sec, 72°C 30sec and 72°C 3min. PCR products were then separated in 1.2% agarose gel running with ladder (1 Kb Plus and 100bp Plus DNA Ladders, Invitrogen). The DNA band at expected size of the circle was extracted by the Qiagen gel purification kit and then cloned into pCR4 Blunt-TOPO vector (Invitrogen) for sequencing.

Plasmids were sequenced and blasted to the JFH-1 sequence. A few vcircRNAs were then validated, containing one or iterated vcircRNA sequences with two identical repeated junction sites flanked at both ends (Supplementary Table 3).

### RNase R treatment and Northern blot analysis

RNase R treatment was performed by incubating 15µg of total RNA with 2.5ul RNaseR (10 U/μl, Abcam) at 37°C for 30 min followed by cleanup using RNA concentrate and clean kit (ZYMO lnc.). For Northern blot analysis of HCV genomic RNA and vcircRNA, 10μg of undigested total RNA samples and the above-mentioned RNase R-treated RNA were added with RNA loading dye (NEB) and denatured for 10min at 70°C. Subsequently, RNA samples were separated in 1% agarose gel containing 1X MOPS-EDTA-Sodium acetate (MESA, Sigma) and 6% formaldehyde, and then transferred onto Zeta-probe membranes (Biorad) by upload capillary blotting overnight. Detection of HCV RNA (HCV nucleotides 84–374) and vcircRNA873/1151 (oligonucleotide probe 5’- CAGGACAACAGGGCCAGCAAGGTGGCGGACATCACAACCA-3’ against the junction) were performed using γ-^32^P-dATP-end labeled (PNK kinase, NEB) probes. Autoradiographs of phosphor screen were quantitated using ImageQuant (GE Healthcare).

### HCV vcircRNA and viral mRNA quantification by qPCR and Droplet digital PCR (ddPCR)

1μg RNA was used to prepare cDNA using the High-Capacity RNA-to-cDNA Kit (Applied Biosystems). qPCR was performed using the PowerUP SYBR green PCR mix in a 96-well format in a CFX96 qPCR machine according to the manufacturer’s instructions (BioRad). HCV RNA abundance at Core, Envelope 1, NS5B regions and vcircRNA∼870/∼1151 was quantified by using primers listed in Supplementary Table 4. Droplet Digital PCR was performed using the Bio-Rad’s QX200 Droplet Digital PCR system (Stanford Genomics). Briefly, 1μg RNA was reverse transcribed using the Superscript III kit and a reverse primer for cluster II vcircRNA (Supplementary Table 4). Subsequently cDNA was mixed with 10μL of QX200ddPCR EvaGreen supermix and 125nM primers. The reaction was then dispensed into sample wells in the DG8 cartridge so that droplets were generated according to the manufacturer’s instruction. The droplets were then transferred to a 96 well plate and run on a thermal cycler with the following conditions: 95°C for 5min, 42 cycles at 98°C for 30sec and 59°C for 1min, 4°C for 5min and 90°C for 5min. After the PCR was completed, the plate was transferred in to a QX200 droplet reader and analyzed by QuantaSoft Analysis Pro software (Bio-Rad). Both HCV RNA in the Core region and cluster II vcircRNA were quantitated separately for each sample, with at least 3 biological repeats.

### Plasmid over-expression of vcircRNA873/1151

The plasmid placcase2-splitGFP was a generous gift from Dr. Jeremy E. Wilusz (Baylor College, Texas). To insert the sequence (5’-3’) of HCV circRNA873/1151 in between the two laccase2 inverted repeats, the plasmid was amplified with the following primer pair to exclude the entire split-GFP-EMCV-IRES region (Figure S 2a) forward 5’- GTAAGTATTCAAAATTCCAAAAT-3’, reverse 5’- CTGCataaaataaaaaaaCTT-3’. The sequence (from 5’ breakpoint to 3’ breakpoint) of vcircRNA873/1151 was amplified from HCV JFH-1 infected cells by using the following primer pair: forward 5’- TTTTATTTTATGCAGTTGCTGGCCCTGTTGTCCT-3’, reverse 5’- ATTTTGAATACTTACGGTGGCGGACATCACAACC-3’. These two PCR products were subjected to InFusion Cloning (Takara Bio.) to generate the overexpression plasmid pc873/1151. For transfection, 1.7×10^6^ 293FT cells were seeded in a 6cm dish and transfected with 3µg pc873/1151. Total RNA was extracted at 2 days post transfection for Northern blot analysis.

### Construction of placasse2-split-GFP plasmids containing IRES sequences from various vcircRNAs

To replace the EMCV-IRES in placcase2-splitGFP with the sequence (5’-3’) of HCV circRNA10/405, two fragments were amplified from placcase2-splitGFP with primer pair 1 and 2. Primer sequences were as followed: pair 1 forward: 5’- CTTGGTACCGAGCTCGGATCCACTAGTCCAGTGTGGTGGAAT-3’, pair 1 reverse: 5’ TTATTAATTAAGTCGACTGCAGAATTCAGATCC-3’; pair 2 forward: 5’-GT CCTGCAGG-GT ATGGTGAGCAAGGGCGAGGAGCT-3’, pair 2 reverse: 5’- AACGGGCCCTCTAGACTCGAGCGGCCGCCTGAGGTGCCAC-3’. A third fragment was amplified from HCV JFH-1 infected cells from 10-405nt. The following primers were used to also include two restriction enzyme sites *PacI* and *SbfI* at its 5’ or 3’ end, respectively: forward 5’- gcagtcgac TTAATTAATAATAGGGGCGACACTCCGCC, reverse 5’- ctcaccatACCCTGCAGGACGTCTTCTGGGCGACGGTTG-3’. The plasmid placcase2-splitGFP was then digested with *BamHI* and *XhoI and* subjected to InFusion Cloning with three fragments mentioned above. The resulting plasmid placcase-splitGFP-10/405 was later digested with *PacI* and *SbfI* and then used in the InFusion Cloning to generate plasmids inserted with the individual sequences (5’-3’) of HCV circRNA30/372, circRNA14/362, circRNA26/369 and circ873/1151 (primers listed in Supplementary Table 4). For transfection, 4.5×10^5^ Huh7 cells per well were seeded in 6-well plates. On the following day, 1.5 μg of each plasmid was transfected using lipofectamine 3000 according to manufacturer’s protocol. At 2 days post transfection, GFP fluorescence images were taken using an inverted microscope Keyence BZ-X700 with BZ-X analyzer software.

### Generation of mutant HCV JFH-1 DNA infectious clones containing a HiBiT tag inserted in the 5’ non-coding region

pHCV-JFH-1 genotype 2a DNA infectious clone was a generous gift from Dr. Charles Rice (Rockefeller University). To insert a HiBiT tag (VSGWRLFKKIS) and U32G mutation to eliminate a stop codon, an upstream fragment of 125bp was amplified from the wildtype pJFH-1 plasmid by nested PCR with the forward primer 5’-AAAAGCAGGCTACTCGAATTCTAATACGACT-3’ and the first reverse primer 5’- AACAGCCGCCAGCCGCTCACGGGAGTGATTCCTGGCGGAGT-3’, and then the second reverse primer 5’-agTTAGCTAATCTTCTTGAACAGCCGCCAGCCGCT-3’. A downstream fragment of 2951bp was amplified with the forward primer 5’- TTCAAGAAGATTAGCTAActgtgaggaactactgtcttc-3’ and the reverse primer 5’- CACGCGATGCCATCGCGGCC-3’. These two fragments were then included in the InFusion Cloning, together with the pJFH-1 plasmid digested with *NotI* and *EcoRI*. The resulting infectious clone was named pHCV-c10/405-Hibit. To generate pHCV-c25/1277-Hibit, an upstream fragment of 139bp was synthesized (AAAAGCAGGCTACTCGAATTCTAATACGACTCACTATAGACCTGCCCCTAATAGGGGCGACACTCCGCCATGAATCACTCCCCGTGAGCGGCTGGCGGCTGTTCAAGAAGATTAGCTGTGAGGAACTACTGTCTTC) by IDT (Philadelphia, USA). The downstream fragment was amplified from the wildtype pJFH-1 plasmid by using forward primer 5’ - TGTGAGGAACTACTGTCTTCACGCAG-3’ and the reverse primer 5’- CACGCGATGCCATCGCGGCC-3’. Both fragments were fused to pJFH-1 plasmid in a similar way as mentioned above.

### Generation of Nanoluciferase-containing HCV DNA infectious clones with GDD (wild-type) or GND mutation in NS5B gene

To construct pHCV-Nluc-WT, the gBlock fragment (19aaCore-Nluc-P2AUbi, Supplementary Table 4) was synthesized, containing a Nanoluciferase gene fused with the first 19 amino acids of HCV JFH-1 Core protein at its N terminus, and a 19-residue porcine teschovirus-1 2A self-cleaving peptide and a Ubi sequence at its C terminus. This fragment was inserted immediately after HCV 5’ UTR, followed with the complete HCV polyprotein. GDD to GND mutation was further introduced in HCV-Nluc-GND by PCR primers (Supplementary Table 4).

### *In-vitro* synthesis of RNA and transfection

Plasmids pHCV-WT (wild-type strain JFH-1), pHCV-c10/405-Hibit, pHCV-c25/1277-Hibit, pHCV- Nluc-WT and pHCV-Nluc-GND were linearized with *XbaI* and transcribed using the T7 MEGAscript kit (Ambion), using the manufacturer’s instructions. PCR products of different lengths (1-226nt-Hibit, 1- 444nt-Hibit and 1-820nt-Hibit) were amplified from pHCV-c10/405- Hibit (primers listed in Supplementary Table 4), and subsequently used as DNA templates for *in vitro* transcribed control RNAs. To detect HiBiT expression from virus mutants, Huh7 cells in 6- well plates were transfected with 3 μg of each *in vitro* transcribed RNA using the TransIT mRNA transfection kit (Mirus Bio) as previously described (33). After 6, 12 or 24 hours of incubation, cells were harvested for HiBiT lytic assay (Promega).

### Small interfering RNA experiments

Two siRNAs against the junction site of vcircRNA were used to deplete vcircRNA873/1151: sense siRNA-a, 5’-CGCCACCUUGCUGGCCCUGUUdTdT-3’ and sense siRNA-b, 5’- GCCACCUUGCUGGCCCUGUUGdTdT-3’. In addition, the following siRNAs against junction sequence of certain IRES-containing circRNAs were synthesized: for depletion of vcircRNA10/405, sense siRNA-a, 5’-GAAGACGUUAAUAGGGGCGACdTdT-3’ and sense siRNA-b, 5’- UCGCCCAGAAGACGUUAAUAGdTdT-3’; for vcircRNA26/369, sense siRNA-a, 5’- AAACCUCAAAGAAACCGCCAUdTdT-3’ and sense siRNA-b, 5’-ACCUCAAAGAAACCGCCAUGAdTdT-3’; for vcircRNA30/372, sense siRNA-a, 5’- AAAGAAAAACCAUGAAUCACUdTdT-3’, sense siRNA-b, 5’- GAAAAACCAUGAAUCACUCCCdTdT-3’ and sense siRNA-c, 5’- AGAAAAACCAUGAAUCACUCCdTdT-3’; for vcircRNA25/1277, sense siRNA 5’ CCCUGGCACCAUCCGCCAUGAdTdT-3’, antisense siRNA 5’-UCAUGGCGGAUGGUGCCAGGGdTdT-3’. The siRNA duplexes were formed by combining sense and their corresponding antisense strands in 1X siRNA Buffer (Dharmacon) as described previously (16). As a negative control siRNA, the following oligonucleotides were used: sense 5′- GAUCAUACGUGCGAUCAGAdTdT-3’ and antisense 5’- UCUGAUCGCACGUAUGAUCdTdT-3’. To knock down HiBiT-tagged vcircRNAs, 3×10^5^ Huh7 cells were seeded overnight in 6-well plates and then transfected with pooled siRNAs mentioned above targeting different IRES-containing vcircRNAs at a final concentration of 100 nM. After incubation for 24 hours, cells were transfected with 3 μg IVT RNAs of HCV-c10/405- Hibit and HCV-c25/1277-HiBiT. Further after 24 hours, HiBiT expression was measured by the HiBiT lytic assay (Promega). To test off-target effects of siRNAs, Huh7 cells were similarly transfected with 100 nM of control siRNA, or siRNAs against cluster I vcircRNAs or cluster II vcirc 873/1151 in 12-well plates. After 24 hours, 1 μg *in vitro* transcribed HCV-Nluc-GND RNA was introduced into cells. After additional 20 hours, HCV Nluc was measured by NanoGlo assay (Promega). To determine the effects of vcircRNAs on HCV replication, 100 nM of siRNA duplexes were transfected into Huh7 cells in 12-well plate using Dharmafect I (Dharmacon) and incubated overnight. Cells were then either infected with JFH-1 at a MOI of 0.5 or transfected with 1.5 μg *in vitro* transcribed wild-type HCV RNA. 48-hour later, total RNA was extracted and vcircRNA∼870/∼1151 and HCV viral RNA levels were assessed by real-time qPCR (Primers listed in Supplementary Table 4).

### Overexpression of polypeptide translated from vcRNA10/405 ORF

To construct overexpression plasmids for the translational product of vc10/405, two gBlock fragments were synthesized by IDT, which contain a HiBiT tag and a linker (GSSG) either N- terminally or C-terminally fused to c10/405 ORF. These two fragments were fused with pcDNA3.1(+) vector by InFusion cloning to generate pC-hibit-10/405ORF and pN-hibit- 10/405ORF. In addition, a third plasmid pSTOP-N-hibit-10/405ORF was constructed as a negative control in which two stop codons were introduced immediately after AUG. For transfection, 8×10^4^ Huh7 cells were seeded in 24-well plate. On the next day, 200ng of IVT RNA of HCV-Nluc-WT were transfected using the TransIT mRNA transfection kit (Mirus Bio).

After 24-hour incubation at 37°C, cells were transfected with 400ng of each plasmid DNA using Lipofectamine 3000 and P3000 Reagent (Invitrogen) according to manufacturer’s instruction.

After additional 24 hours, Nluc activity from HCV was measured by the NanoGlo assay.

### Statistical analysis

Ordinary one-way ANOVA or two-way ANOVA were used to evaluate the data for statistical differences (*, p < 0.05, **p<0.01, ***p<0.001, ****p<0.0001) in GraphPad Prism (version 9.0).

## Funding

Funding for this project was provided by grants from the National Institutes of Health (R01 AI069000, R21 AI151715) and from the Chan Zuckerberg BioHub.

## Author contributions

P.S., Q.C., P.B., T.C. and K.S. conceived and designed the study. Q.C., P.B. and T.C. conducted the experiments. P.S and Q.C. wrote the manuscript. S.L. performed the graphic analyses of the vcircRNAs.

## Competing interests

The authors declare that they have no competing interests.

## Data and materials availability

All data are available in the main text and the Supplementary materials. GEO accession will be available to public before the paper is accepted for publication.

**Supplementary Figures S1-S6 are attached.**

**Supplementary Tables in Excel forms are available for this paper:**

Supplementary Table 1. Novel vcircRNA junction reads detected in infected cells with or without RNase R treatment.

Supplementary Table 2. Detection of vcircRNA junction reads in RNase R-treated samples by CIRI.

Supplementary Table 3. Sanger sequencing of RT-PCR products that contain a complete copy of indicated viral circRNA sequence flanked by two identical junction sites.

Supplementary Table 4. List of primers used in the study.

## Acknowledgements

We acknowledge Dr. Julia Salzman for many helpful comments during this study. We are also very grateful to Dr. Karla Kirkegaard for critical reading the manuscript.

## Notes

### Competing Interest Statement

The authors have declared no competing interest.

