## Supplementary material for "Virus-derived circular RNAs populate hepatitis C virus-infected cells": Suppl. Fig

Figure S1

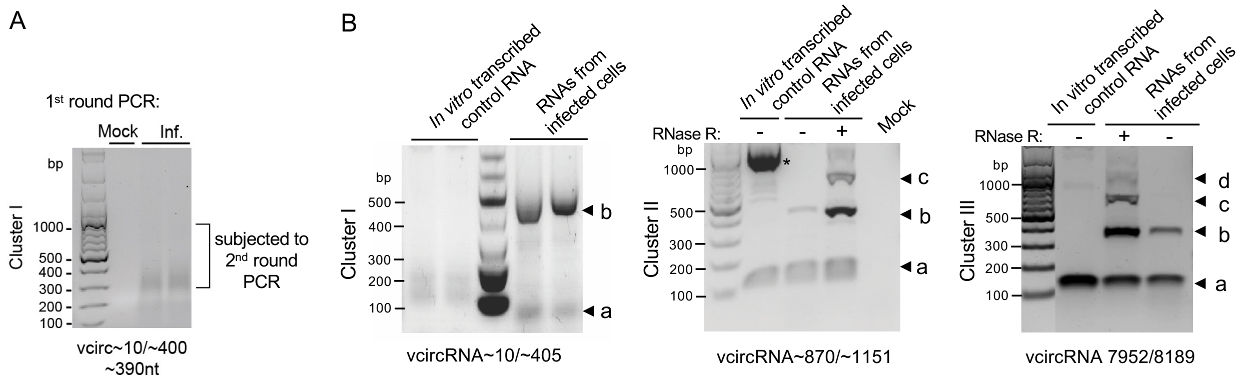


**Figure S1. Viral circRNAs are detected from virus-infected cells but not from *in vitro* transcribed HCV RNA, related to Figure 2.** (A) The first round of nested PCR for amplification of cluster I vcircRNAs. Products in the indicated region was gel-purified and subjected to the second round of PCR. (B) PCR amplification of vcircRNAs from *in vitro*-transcribed full-length HCV control RNA or total RNAs isolated from HCV-infected cells. Divergent primers that flank or span the junction of each cluster of vcircRNAs were used. Products from cluster I (left panel, 2nd round PCR is shown), cluster II (middle panel) and cluster III (right panel) are shown as b, c, d, indicating one or more copies of the vcircRNA sequence are amplified. * in the middle panel indicates an unspecific PCR band from *in vitro*-transcribed HCV RNA. It was purified, sequenced and proved not to have discontinuous junction sites of vcircRNAs (data available upon request).

Figure S2

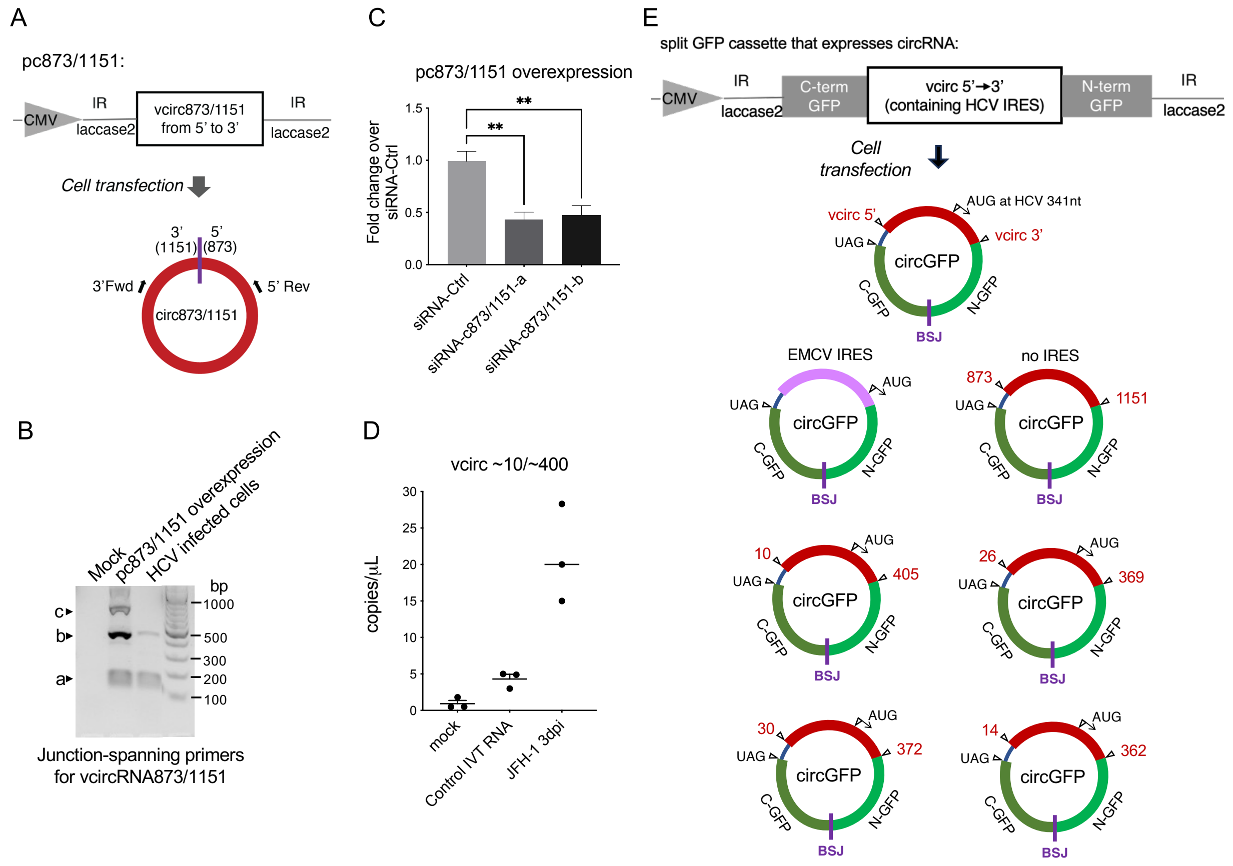


**Figure S2. Overexpression of cluster II vcircRNA 873/1151 (related to Figure 2) and translation of circular GFP from split-GFP plasmids containing HCV IRES from cluster I vcircRNAs (related to Figure 3).** (A) Schematic view of the plasmid over-expressing circRNA 873/1151. A previously reported plasmid system[1] was used in which intronic inverted repeats (IR) from the *Drosophila laccase2* gene facilitate the back-splicing of the flanked RNA upon transcription, hence generating a circular RNA. After transfection, RT-PCR was performed to verify the exogenous expression of vcircRNA using divergent primers flanking the junction 873/1151, and RT-PCR products were displayed on an 1.2% agarose gel shown in (B). (C) Depletion of circ873/1151. Huh7 cells were transfected with each of two siRNA targeting the junction sequence of vcirc873/1151 at a final concentration of 100nM, and subsequently transfected with plasmid pc873/1151. Circ873/1151 abundance was quantified by real-time PCR using primers spanning the junction. Data are represented as means ± SEM. Statistical significance was determined by ordinary one-way ANOVA. **, $p<0.01$. (D) Droplet digital PCR to quantify the IRES-containing cluster I vcircRNAs. Total RNAs were extracted from mock-infected cells and JFH-1-infected cells at three days after infection. Control *in vitro* transcribed full-length HCV RNA was used as a negative control for vcircRNAs formation. (E) Schematic views of a split-GFP plasmid that contains an IRES sequence from selected cluster I vcircRNAs. The plasmid cassette has the GFP coding sequence separated by the indicated vcircRNA sequence from 5' to 3' breakpoints. Upon transcription, the flanking *laccase2* inverted repeats drive the formation of a circRNA that juxtaposes the N- and C-terminus of the GFP ORF. Translation of the GFP ORF is then initiated by the IRES within the circular GFP RNA. Split-GFP plasmids containing individual cluster I vcircRNA are shown here.

### Figure S3

**Predicted vcirc10/405 ORF is underlined** (10/405 full sequence is shown below with 3' sequence in blue and 5' in green):

UCCCCUGUGAGGAACUACUGUCUUCACGCAGAAAGCGCCUAGCCAUGGCGUUAGUAUGAGUGUCGUACAGCCUCCAGGC
CCCCCCCUCCCGGGAGAGCCAUAGUGGUCUGCGGAACCGGUGAGUACACCGGAAUUGCCGGGAAGACUGGGUCCUUUCU
UGGAUAAACCCACUCUAUGCCCGGCCAUUUGGGCGUGCCCCCGCAAGACUGCUAGCCGAGUAGCGUUGGGUUGCGAAAG
GCCUUGUGGUACUGCCUGAUAGGGCGCUUGCGAGUGCCCCGGGAGGUCUCGUAGACCGUGCACCA$_{341}$UGAGCACAAAUC
CUAAACCUCAAAGAAAAACCAAAAGAAACACCAACCGUCGCCCAGAAGACGU|UAAUGGCGACACUCCGCCAU$_{33}$GAAUCAC

405|10
Junction site

**Predicted vcirc10/405 protein sequence (29aa):**

MSTNPKPQRKTKRNTNRRPEDV|NRGDTPP.

**Predicted ORF of HiBiT-tagged vcirc10/405 from HCV-c10/405-Hibit** (HiBiT sequence in pink):

A$_{341}$UGAGCACAAAUCCUAAACCUCAAAGAAAAACCAAAAGAAACACCAACCGUCGCCCAGAAGACGU|UAAUGGCGACACUCCG
CCAG$_{33}$GAAUCACUCCC$_{43}$GUGAGCGGCUGGCGGCUGUUCAAGAAGAUUAGCUAA

**Predicted vcirc10/405 protein sequence with in-frame HiBiT at C terminus:**

MSTNPKPQRKTKRNTNRRPEDV|NRGDTPPGITPVSGWRLFKKIS.

**Potential ORF predicted from vcirc25/1277** (3' sequence in blue and 5' in green):

a$_{341}$tgagcacaaatcctaaacctcaaagaaaaaccaaaagaaacaccaaccgtcgcccagaagacgttaagttcccgggcggcggccagatcgttggcggagtatacttgttgc
cgcgcaggggccccaggttgggtgtgcgcacgacaaggaaaacttcggagcggtcccagccacgtgggagacgccagcccatccccaaagatcggcgctccactggcaaggc
ctggggaaaaccaggtcgcccctggcccctatatgggaatgagggactcggctgggcaggatggctcctgtcccccgaggctctcgcccctcctggggccccactgacccccggc
ataggtcgcgcaacgtgggtaaagtcatcgacaccctaacgtgtggctttgccgacctcatggggtacatccccgtcgtaggcgccccgcttagtggcgccgccagagctgtcgcgca
cggcgtgagagtcctggaggacggggttaattatgcaacagggaacctacccggtttcccttttctatcttcttgctggccctgttgtcctgcatcaccgttccggtctctgctgcccaggtg
aagaataccagtagcagctacatggtgaccaatgactgctccaatgacagcatcacttggcagctcgaggctgcggttctccacgtccccgggtgcgtcccgtgcgagagagtgggg
aatacgtcacggtgttgggtgccagtctcgccaaacatggctgtgcggcagcccggtgccctcacgcagggtctgcggacgcacatcgatatggttgtgatgtccgccaccttctgctct
gctctctacgtgggggacctctgtggcggggtgatgctcgcggcccaggtgttcatcgtctcgccgcagtaccactggtttgtgcaagaatgcaattgctccatctaccctggcacca|tcc
gccatgaatcactcccc$_{44}$tgtgaggaactactgtcttcacgcagaaagcgcctagccatggcgttagtat$_{96}$ga

1277|25
Junction site

**Predicted vcirc25/1277 protein sequence (336aa):**

MSTNPKPQRKTKRNTNRRPEDVKFPGGGQIVGGVYLLPRRGPRLGVRTTRKTSERSQPRGRRQPIPKDRRSTGKAWGKPGRPWPL
YGNEGLGWAGWLLSPRGSRPSWGPTDPRHRSRNVGKVIDTLTCGFADLMGYIPVVGAPLSGAARAVAHGVRVLEDGVNYATGNLP
GFPFSIFLLALLSCITVPVSAAQVKNTSSSYMVTNDCSNDSITWQLEAAVLHVPGCVPCERVGNTSRCWVPVSPNMAVRQPGALT
QGLRTHIDMVVMSATFCSALYVGDLCGGVMLAAQVFIVSPQYHWFVQECNCSIYPGTI|RHESLPCEELLSSRRKRLAMALV*

**Predicted ORF of HiBiT-tagged vcirc25/1277 from HCV-c25/1277-Hibit** (HiBiT sequence in pink):

a$_{341}$tgagcacaaatcctaaacctcaaagaaaaaccaaaagaaacaccaaccgtcgcccagaagacgttaagttcccgggcggcggccagatcgttggcggagtatacttgttgcc
gcgcaggggccccaggttgggtgtgcgcacgacaaggaaaacttcggagcggtcccagccacgtgggagacgccagcccatccccaaagatcggcgctccactggcaaggcct
ggggaaaaccaggtcgcccctggcccctatatgggaatgagggactcggctgggcaggatggctcctgtcccccgaggctctcgcccctcctggggccccactgacccccggcat
aggtcgcgcaacgtgggtaaagtcatcgacaccctaacgtgtggctttgccgacctcatggggtacatccccgtcgtaggcgccccgcttagtggcgccgccagagctgtcgcgcacg
gcgtgagagtcctggaggacggggttaattatgcaacagggaacctacccggtttcccttttctatcttcttgctggccctgttgtcctgcatcaccgttccggtctctgctgcccaggtgaa
gaataccagtagcagctacatggtgaccaatgactgctccaatgacagcatcacttggcagctcgaggctgcggttctccacgtccccgggtgcgtcccgtgcgagagagtggggaat
acgtcacggtgttgggtgccagtctcgccaaacatggctgtgcggcagcccggtgccctcacgcagggtctgcggacgcacatcgatatggttgtgatgtccgccaccttctgctctgctc
tctacgtgggggacctctgtggcggggtgatgctcgcggcccaggtgttcatcgtctcgccgcagtaccactggtttgtgcaagaatgcaattgctccatctaccctggcacca|tccgcca
tgaatcactcccc$_{44}$GTGAGCGGCTGGCGGCTGTTCAAGAAGATTAGCtgtgaggaactactgtcttcacgcagaaagcgcctagccatggcgttagtat$_{96}$ga

**Predicted vcirc25/1277 protein sequence inserted with an in-frame HiBiT:**

MSTNPKPQRKTKRNTNRRPEDVKFPGGGQIVGGVYLLPRRGPRLGVRTTRKTSERSQPRGRRQPIPKDRRSTGKAWGKPGRPWPL
YGNEGLGWAGWLLSPRGSRPSWGPTDPRHRSRNVGKVIDTLTCGFADLMGYIPVVGAPLSGAARAVAHGVRVLEDGVNYATGNLP
GFPFSIFLLALLSCITVPVSAAQVKNTSSSYMVTNDCSNDSITWQLEAAVLHVPGCVPCERVGNTSRCWVPVSPNMAVRQPGALT
QGLRTHIDMVVMSATFCSALYVGDLCGGVMLAAQVFIVSPQYHWFVQECNCSIYPGTI|RHESLPVSGWRLFKKISCEELLSSRR
KRLAMALV*

**Figure S3. Predicted opening reading frames (ORFs) and protein sequences of vcirc10/405 and vcirc25/1277 derived from wild-type HCV, and HiBiT-tagged vcirc10/405 and vcirc25/1277 from HCV mutants, related to Figure 3.**

Figure S4

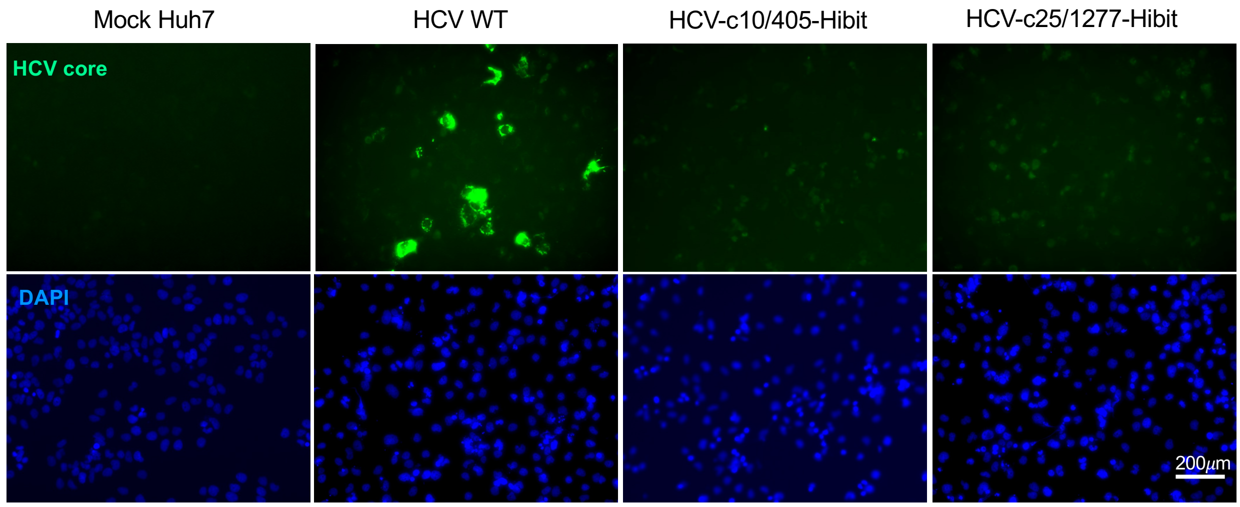


**Figure S4. Measurement of growth phenotypes of HiBiT-inserted HCV mutants, related to Figure 3.** Huh7 cells were transfected with full-length viral RNAs generated from plasmid wildtype[2], pHCV-c10/405-Hibit and pHCV-c25/1277-Hibit. At three days post transfection, immunofluorescent assay was performed by using anti-HCV core antibody at 1:2000.

Figure S5

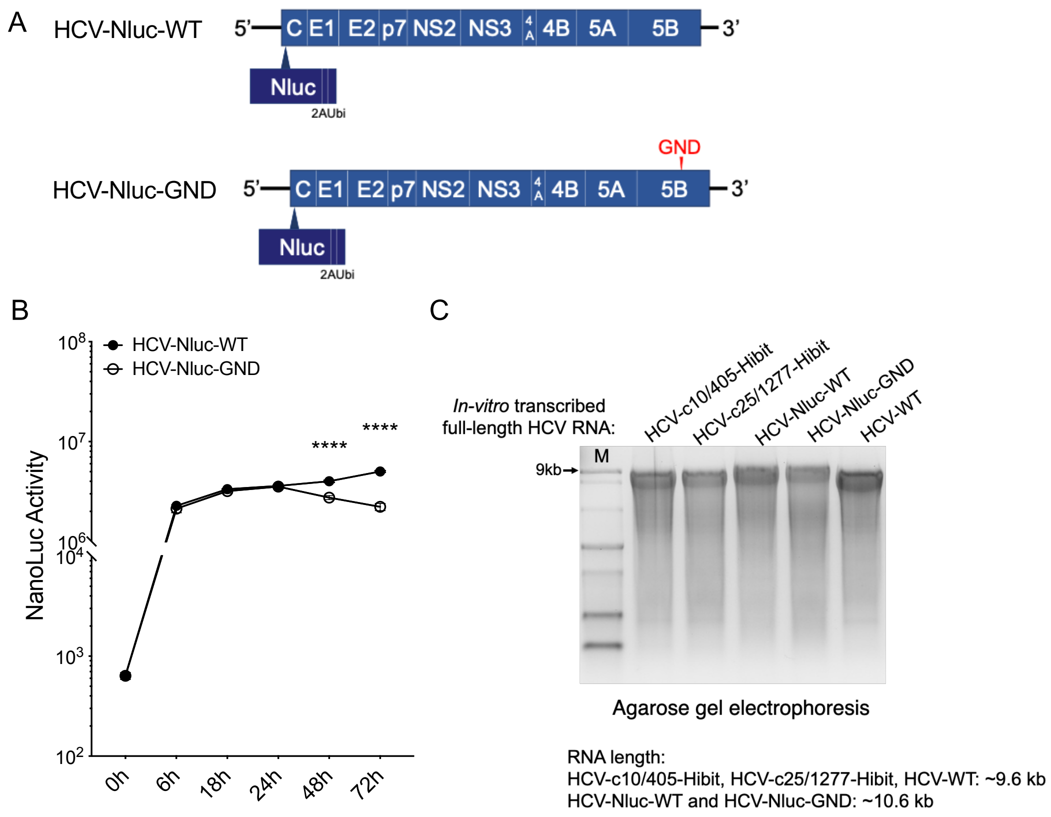


**Figure S5. Construction of replication-competent and -deficient HCV JFH-1 infectious clones carrying a Nanoluciferase insertion, related to Figure 5**. (A) Comparison of HCV-Nluc-WT and HCV-Nluc-GND, the latter contains a GDD-to-GND mutation in NS5B, generating a non-functional viral polymerase. 2A/Ubi promotes "stop-carry on" translational recoding[3]. (B) Kinetics of nanoluciferase expression after transfection of *in vitro*-transcribed HCV-Nluc-WT and HCV-Nluc-GND RNAs into Huh7 cells shows that both RNAs can be translated. Data are shown as means ± SEM. Statistical significance was determined by ordinary two-way ANOVA. ****$p<0.0001$. (C) *In vitro* transcribed full-length HCV RNAs from individual plasmids of HCV mutants were examined for their integrity and lengths by agarose gel electrophoresis.

#### Figure S6

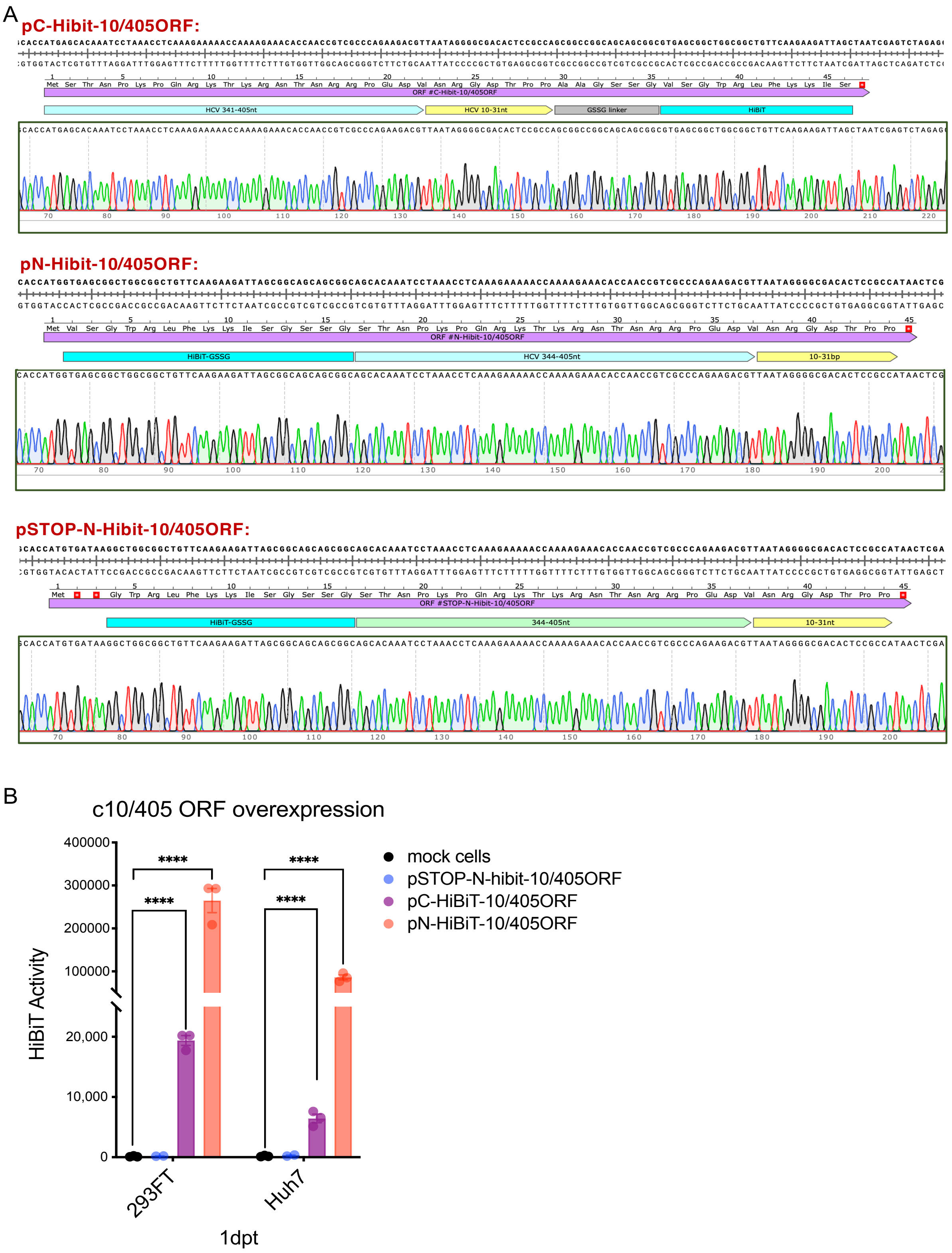
A
pC-Hibit-10/405ORF:
ORF #C-Hibit-10/405ORF
HCV 341-405nt
HCV 10-31nt
GSSG linker
HiBiT
pN-Hibit-10/405ORF:
ORF #N-Hibit-10/405ORF
HiBiT-GSSG
HCV 344-405nt
10-31bp
pSTOP-N-Hibit-10/405ORF:
ORF #STOP-N-Hibit-10/405ORF
HiBiT-GSSG
344-405nt
10-31nt
B
c10/405 ORF overexpression
HiBiT Activity
400000
300000
200000
100000
20,000
10,000
0
****
****
****
****
293FT
Huh7
1dpt
mock cells
pSTOP-N-hibit-10/405ORF
pC-HiBiT-10/405ORF
pN-HiBiT-10/405ORF

**Figure S6. Over-expression of the polypeptide from cluster I vcircRNA10/405 opening reading frame, related to Figure 5**. (A) Schematic view of plasmids pC-Hibit-10/405ORF and pN-Hibit-10/405ORF that contain either a C- or N-terminal HiBiT tag in the vcircRNA10/405 ORF. A third control plasmid pSTOP-N-Hibit-10/405 contains two immediate stop codons placed after the first amino acid of the N-terminal HiBiT tagged ORF. (B) 1 μg of each plasmid was transfected into either 293FT or Huh7 cells in 12-well plates. HiBiT luciferase activity was measured at 24 hours post transfection by HiBiT Lytic Assay (Promega). Data are shown as means ± SEM. Statistical significance was determined by multiple t-test between indicated groups. ****$p<0.0001$.
